# Ifit1 regulates norovirus infection and enhances the interferon response in murine macrophage-like cells

**DOI:** 10.1101/611236

**Authors:** Harriet V. Mears, Edward Emmott, Yasmin Chaudhry, Myra Hosmillo, Ian G. Goodfellow, Trevor R. Sweeney

## Abstract

**Background:** Norovirus, also known as the winter vomiting bug, is the predominant cause of non-bacterial gastroenteritis worldwide. Disease control is predicated on a robust innate immune response during the early stages of infection. Double-stranded RNA intermediates generated during viral genome replication are recognised by host innate immune sensors in the cytoplasm, activating the strongly antiviral interferon gene programme. Ifit proteins, which are highly expressed during the interferon response, have been shown to directly inhibit viral protein synthesis as well as regulate innate immune signalling pathways. Ifit1 is well-characterised to inhibit viral translation by sequestration of eukaryotic initiation factors or by directly binding to the 5’ terminus of foreign RNA, particularly those with non-self cap structures. However, noroviruses have a viral protein, VPg, covalently linked to the 5’ end of the genomic RNA, which acts as a cap substitute to recruit the translation initiation machinery.

**Methods:** Ifit1 knockout RAW264.7 murine macrophage-like cells were generated using CRISPR-Cas9 gene editing. These cells were analysed for their ability to support murine norovirus infection, determined by virus yield, and respond to different immune stimuli, assayed by quantitative PCR. The effect of Ifit proteins on norovirus translation was also tested *in vitro*.

**Results:** Here, we show that VPg-dependent translation is completely refractory Ifit1-mediated translation inhibition *in vitro* and Ifit1 cannot bind the 5’ end of VPg-linked RNA. Nevertheless, knockout of Ifit1 promoted viral replication in murine norovirus infected cells. We then demonstrate that Ifit1 promoted interferon-beta expression following transfection of synthetic double-stranded RNA but had little effect on toll-like receptor 3 and 4 signalling.

**Conclusions:** Ifit1 is an antiviral factor during norovirus infection but cannot directly inhibit viral translation. Instead, Ifit1 stimulates the antiviral state following cytoplasmic RNA sensing, contributing to restriction of norovirus replication.

## Introduction

The *Caliciviridae* family of small positive-sense RNA viruses comprises 11 genera, including *Norovirus* and *Sapovirus*. Noroviruses are the leading cause of non-bacterial gastroenteritis in humans, accounting for 18% of acute gastroenteric disease worldwide (1). While recent advancements in human intestinal organoids have made it possible to study human noroviruses in culture (2), murine norovirus (MNV) remains a valuable model for dissecting interactions between noroviruses and their host, owing to readily cultivable permissive cell lines and a flexible reverse genetics system (3).

The innate immune response to viral infection is essential for the control of norovirus replication and clearance (4). Sensing of calicivirus infection is predominantly mediated by cytoplasmic double-stranded RNA sensors; both RIG-I and MDA5 have been implicated in controlling the innate immune response at different stages of infection (5–7). By contrast TLR3, an endosomal dsRNA sensor, has little effect on norovirus replication (5). RIG-I and MDA5 signalling converge on the activation of the antiviral signalling complex MAVS, which recruits TBK1 to induce the phosphorylation of interferon regulatory factor (IRF) 3. Activated IRF-3 dimerises and translocates into the nucleus where it promotes the transcription of type I interferon (IFN) and early antiviral genes.

During the antiviral response, among the strongest upregulated IFN stimulated genes are the IFIT family of RNA binding proteins (8–10). In humans, IFIT1 directly inhibits the translation of non-self RNAs at the initiation stage, by binding over the 5’ terminus, occluding the recruitment of eukaryotic translation initiation factor (eIF) 4F (11–13). IFIT1 binding is highly specific for capped mRNA which lacks methylation at the first or second cap-proximal nucleotides (cap0) (14). Murine Ifit1 similarly binds cap0 RNA and mediates the inhibition of cap0 viruses *in vivo* (11, 15–18). It is important to note, however, that murine Ifit1 and human IFIT1, which share 52% sequence identity, have distinct evolutionary origins, with murine Ifit1 being more closely related to another gene family member, IFIT1B (19).

However, IFIT1 may have antiviral activity independent of its RNA binding capability. IFIT1 was reported to inhibit hepatitis C virus replication (20, 21), by binding to eIF3 to prevent viral translation initiation (22–24). Additionally, direct binding to the human papilloma virus DNA helicase, E1, was reported to inhibit viral DNA replication (25, 26). IFIT1 also modulates different stages of the host innate immune response during both viral and bacterial infection (27–29) and may regulate the inflammatory response in human astrocytes (30). MNV can antagonise innate immune sensing and was consequently shown to inhibit the expression of a number of interferon-stimulated genes, including Ifit2 (31). However, associations between noroviruses and other members of the Ifit family have not been established.

We investigated whether Ifit1 played a role in the antiviral response to calicivirus infection. We show that Ifit1 knockout promoted MNV replication in a macrophage cell line. However, calicivirus translation was not inhibited by Ifit1. Instead, we show that Ifit1 knockout cells have impaired cytoplasmic double-stranded RNA sensing, resulting in a weaker type I IFN response, which permits increased viral replication.

## Materials and Methods

### Cells viruses and plasmids

Murine macrophage RAW264.7, microglial BV2 and Crandell-Rees feline kidney cells were cultured in Dulbecco’s modified Eagle’s medium (DMEM) with 10% (v/v) foetal calf serum (FCS) and 1% penicillin/streptomycin (P/S). LLC-PK cells, expressing bovine viral diarrhoea virus NPro to render them IFN-deficient, were cultured in Eagle’s minimal essential medium (EMEM) supplemented with 200 μM glycochenodeoxycholic acid (GCDCA; Sigma), 2.5% FCS, and 1% P/S (32). MNV-1 strain CW.1 was recovered from the pT7:MNV-G 3’Rz plasmid as described (33). Feline calicivirus (FCV) strain Urbana was recovered from the pQ14 full length infectious clone (34). The porcine sapovirus (PSaV) Cowden tissue culture adapted strain was obtained from K. O. Chang (Kansas State University) and recovered from the full-length infectious clone pCV4A (35). For lentivirus generation, psPAX2 (Addgene plasmid # 12260) and pMD2.G (Addgene plasmid # 12259) were gifts from Didier Trono. For bacterial expression, murine Ifit1 (NM_008331.3), Ifit2 (NM_008332.3) and Ifit3 (NM_010501.2) were cloned between NcoI and XhoI sites in pTriEx1.1, to contain a C-terminal His_8_ tag.

### Knockout cells

Five guide RNAs designed against the 5’ end of the second exon of Ifit1 (Table 1), were cloned into lentiCRISPR v2 (36). 3 μg guide RNA plasmid was cotranfected into 5 × 10^6^ HEK293T cells with 3 μmg psPAX2 packaging vector and 1.5 μg pMD2.G VSV-G envelope vector using lipofectamine 2000 (Invitrogen). Supernatants were harvested over 72 hours, pooled and used directly for transduction of subconfluent RAW264.7 cells. After 3 days, transduced cells were selected with puromycin for one week, before single cell clones were generated by dilution in 96 well plates. Knockout was verified by western blotting after treatment with murine IFNβ for 12 hours.

**Table 1.**
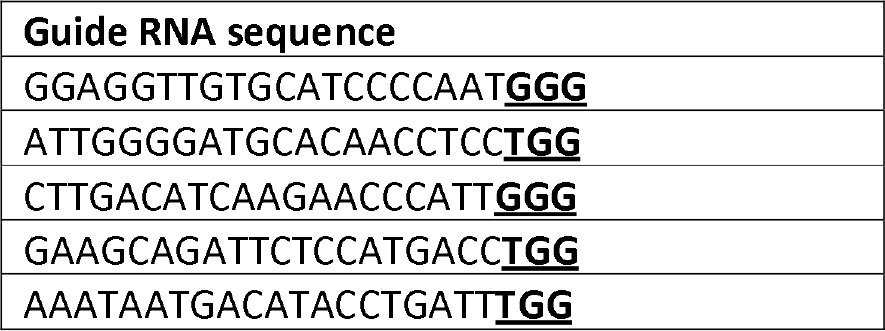
Guide RNA sequences for CRISPR-Cas9 knockout of Ifit1. Guide RNAs were generated using crispr.mit.edu and cloned into LentiCRISPRv2 (36). 3’ protospacer adjacent motif (PAM) sequences are underlined in bold.

### Infections

Cells were infected for 1 hour at 37 °C at the indicated multiplicity of infection (MOI). Cells were harvested by freezing at the indicated times and titres were determined by 50% tissue culture infectious dose (TCID_50_) in BV2 cells, as described (3), performed in technical quadruplicate. Plates were scored by cytopathic effect after 5 days and titres were calculated by the Reed and Muensch method (3).

### Stimulation of RAW264.7 cells

Cells were treated with 10 ng/mL LPS (Sigma) or 1 μg/mL polyI:C (Sigma), or transfected with 2 μg polyI:C using lipofectamine 2000 (Invitrogen). Cells were harvested by washing twice in PBS before lysis in passive lysis buffer (Promega) and RNA was extracted using TRIreagent (Sigma).

### RT-qPCR

For RT-qPCR analysis, cDNA was generated using Moloney murine leukemia virus (M-MLV) reverse transcriptase (Promega) with random hexamer primers. qPCR was performed on cDNA using primers for murine IFNβ (37), TNFα (38) and GAPDH, using SYBR green (Eurogentec). Data were normalised against GAPDH, expressed as fold change over mock (2^−ddCT^).

### Western blotting

Cell lysates were separated in 12.5% SDS-PAGE and transferred to 0.45 μm nitrocellulose membrane by semi-dry blotting. Membranes were blocked in 5% milk phosphate buffered saline with 0.1% tween-20 (PBS-T) and primary antibodies were incubated in 5% BSA PBS-T at 4 °C overnight. Anti-Ifit1 (Santa Cruz, sc-134949, rabbit polyclonal) was used at 1:500, anti-Ifit2/3 (ProteinTech, 12604-1-AP, rabbit polyclonal) was used at 1:800 and anti-GAPDH (Invitrogen, AM4300, mouse monoclonal) was used at 1:8000. Blots were imaged on an Odyssey CLx Imaging System using IRDye secondary antibodies (Li-Cor) at 1:10000 in PBS-T.

### RNA extraction and in *vitro transcription*

Preparation of VPg-linked RNA from MNV, FCV (39, 40) and PSaV (32) infected cells was performed as described using the GenElute total RNA extraction kit (Sigma). *In vitro* transcribed RNAs were generated with T7 polymerase (New England Biosciences) from linearised plasmids and subsequently capped using the ScriptCap Capping System (CellScript).

### Recombinant protein purification

Recombinant Ifit1, Ifit2 and Ifit3 were expressed in BL21 (DE3) Star Escherichia coli (Invitrogen). Cells were grown to an OD600 of ~1.0 in 2× TY media at 37 °C. Expression was induced with 1 mM isopropyl b-D-1-thogalactopyranoside at 22 °C for 16 hours. Cells were harvested in a lysis buffer containing 400 mM KCl, 40 mM Tris pH 7.5, 5% glycerol, 2 mM DTT and 0.5 mM phenylmethylsophonyl fluoride with 1 mg/mL lysozyme. Proteins were purified by affinity chromatography on NiNTA agarose (Qiagen), followed by FPLC on MonoQ (GE Healthcare) as described (41).

### *In vitro* translation

8 nM cap0 or 20 ng/μL VPg-linked RNA was translated using the Flexi Rabbit Reticulocyte Lysate system (Promega) in the presence or absence of 1.5 μM Ifit proteins, including 5 uCi EasyTag™ L-[^35^S]-Methionine (Perkin-Elmer). After 90 min at 30 °C, reactions were terminated by addition of 50 mM EDTA and 0.5 μg/μL RNaseA. Labelled proteins were separated by 12.5% PAGE and detected by autoradiography using an FLA7000 Typhoon Scanner (GE).

### Primer extension inhibition

Primer extension inhibition assays were performed as described (12). Briefly, 1 nM cap0 or VPg-linked RNA were incubated with 1.5 μM Ifit proteins for 10 minutes at 37 °C in reactions containing 20 mM Tris pH 7.5, 100 mM KCl, 2.5 mM MgCl_2_, 1 mM ATP, 0.2 mM GTP, 1 mM DTT and 0.25 mM spermidine. Reverse transcription (RT) was carried out using 2.5 U avian myeloblastosis virus (AMV) reverse transcriptase (Promega) and a ^32^P-labelled primer in the presence of 4 mM MgCl_2_ and 0.5 mM dNTPs. Primer sequences used for RT were CCTGCTCAGGAGGGGTCATG (MNV-1), GTCATAACTGGCACAAGAAGG (FCV) and GTCGTGGGGTGCCAGAAATC (PSaV). Sequencing reactions were performed using the Sequenase Version 2.0 DNA Sequencing Kit (ThermoFisher) in the presence of ^35^S-labelled ATP. cDNA products were resolved on 6% denaturing PAGE and detected by autoradiography using an FLA7000 Typhoon Scanner (GE).

## Results

### Ifit1 inhibits MNV in RAW264.7 cells

We examined the effect of Ifit1 on calicivirus replication, using MNV as a model. Ifit1 knockout RAW264.7 cell lines were generated by CRISPR-Cas9 gene editing and complete knockout was verified by western blotting. Ifit1 expression was undetectable in two independent Ifit1^−/−^ clones after 12 hours treatment with IFNβ, while expression of Ifit2 and Ifit3 was maintained (Figure 1A). Wildtype and Ifit1^−/−^ RAW264.7 cells were then infected with MNV-1 at low or high MOI and samples were harvested by freezing at the indicated time points. Viral titres were determined by TCID_50_ assay in BV2 cells.

**Figure 1.**
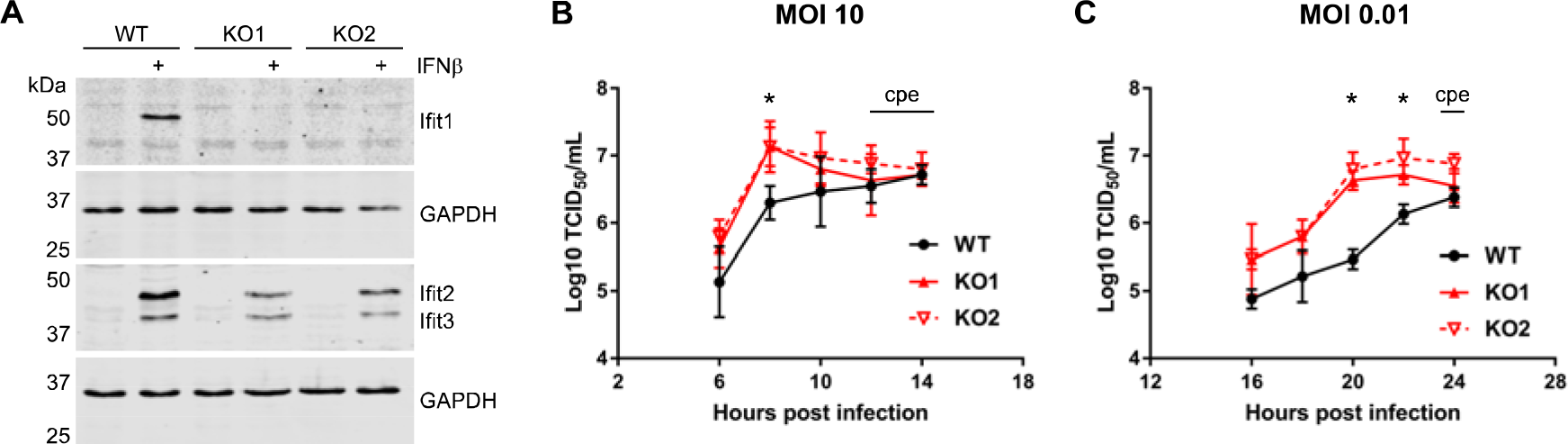
Ifit1 decreases MNV infection in RAW264.7 cells. **A.** Ifit1 knockout RAW264.7 cells were generated by CRISPR-Cas9 gene editing. Cells were stimulated with IFNβ for 12 hours then analysed by western blotting against Ifit1 and Ifit2/Ifit3. GAPDH was included as a loading control for each membrane. **B.C.** Infection of wildtype (WT) and Ifit1 knockout (KO) RAW264.7 cells at **(B)** high or **(C)** low multiplicity of infection (MOI) with murine norovirus (MNV-1). Viral titres were determined by 50% tissue culture infectious dose (TCID_50_) in BV2 cells and expressed as log_10_-transformed values. At late time points, indicated, severe cytopathic effect (cpe) was visible. Graphs show the mean and the standard error of three biological replicates. Titres were compared between WT and KO cells for each time point by two-tailed student’s t-test. Asterisks indicate that a statistically significant difference (p < 0.05) was observed for both KO cell lines.

In Ifit1^−/−^ cells infected at a high multiplicity of infection, MNV-1 titres were slightly higher than wild-type cells at 6-8 hours post infection (Figure 1B). By 12-14 hours post infection viral titres from wildtype and knockout cells were similar. At these times, a high degree of cytopathic effect was observed, hence infection did not progress any further. When infected at low multiplicity, the differences between wildtype and Ifit1^−/−^ cells were more apparent (Figure 1C). Infection of Ifit1^−/−^ cells resulted in up to 20× higher MNV-1 yields compared to wildtype cells over the course of the infection, suggesting that Ifit1 has antiviral activity during norovirus infection.

### Ifit1 cannot inhibit VPg-dependent translation

Ifit1 primarily mediates its antiviral activity by binding to the 5’ cap of non-self RNA, to occlude translation factor recruitment and prevent viral translation. However, members of the *Caliciviridae* family possess a viral protein, VPg, covalently linked to the 5’ end of the genome which promotes viral translation, in place of a 5’ cap (32, 39, 42–44). Since knockout of Ifit1 promoted MNV replication *in vitro*, this suggests that Ifit1 may restrict MNV replication directly, by inhibiting viral translation, or indirectly, by creating a cellular environment which is less permissive to infection. To differentiate these possibilities, we first examined whether Ifit1 could inhibit calicivirus translation *in vitro*. A similar in vitro translation approach was originally used by Guo et al. to describe the activity of IFIT proteins (22), and since has been successfully used to investigate IFIT1 translation inhibition on human parainfluenza virus (45) and Zika virus model RNAs (41).

To generate VPg-linked RNA for examination, total RNA was extracted from cells infected with MNV, PSaV or FCV. For PSaV and FCV, translation from VPg-linked RNA prepared in this way predominantly consists of VP1, the major viral capsid protein, which is translated from a highly abundant subgenomic RNA (Figure 2A) (32, 40). VPg-linked RNAs were translated in rabbit reticulocyte lysate in the presence or absence of recombinantly expressed and purified murine Ifit1, Ifit2 and Ifit3. ^35^S-Met-labelled translation products were separated by SDS-PAGE and detected by autoradiography.

**Figure 2.**
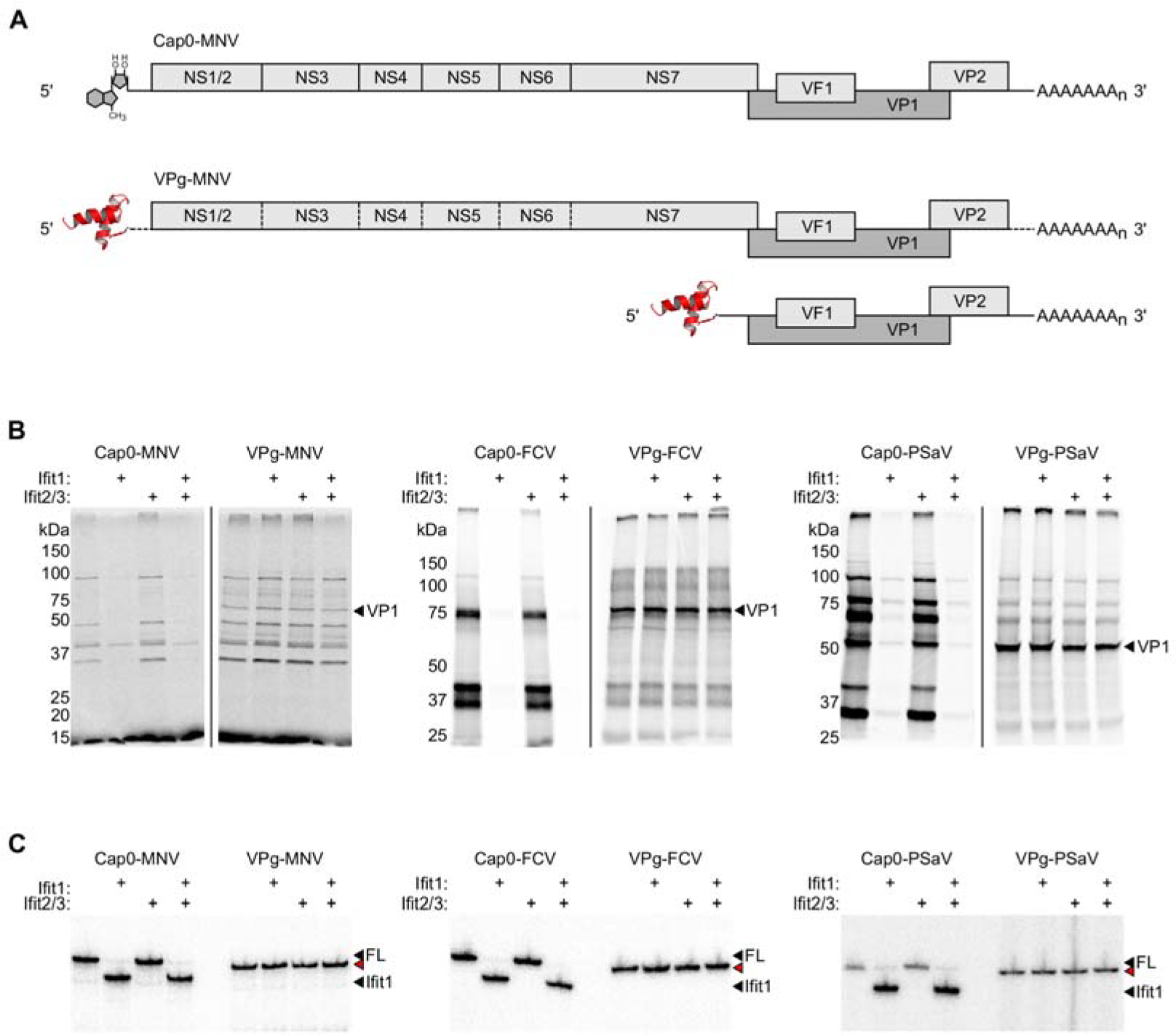
Calicivirus translation is resistant to Ifit1 inhibition. **A.** Schematic representations of *in vitro* transcribed cap0 genomic RNA or VPg-linked genomic and subgenomic RNAs, purified from infected cells, used for *in vitro* translation and toeprint assays. **B.** *In vitro* translation of cap0 or VPg-linked RNA from murine norovirus (MNV), porcine sapovirus (PSaV) and feline calicivirus (FCV). VP1, the dominant protein product produced from the VPg-linked subgenomic RNA, is indicated. **C.** Toeprint analysis of MNV, PSaV and FCV VPg-linked and cap0 RNA. Ifit1 binding is indicated by a cDNA product 6-7 nt shorter than the full-length signal (FL), indicated by black arrowheads. Red arrowheads indicate a 1-2 nt shorter full-length signal on VPg-linked RNAs.

Full-length *in vitro* transcribed cap0 RNA was included as a positive control for Ifit1 activity (Figure 2A). As expected, Ifit1 strongly inhibited the translation of artificial cap0 viral RNA (12), but had no effect on the translation of VPg-linked RNAs (Figure 2B). Addition of Ifit2 and Ifit3 did not enhance translation inhibition on any RNA tested.

Consistently, we observed no evidence of direct Ifit1 binding to VPg-linked RNA when examined in a primer extension inhibition assay, an approach we have used previously to quantify IFIT binding to different RNAs (12, 41). Ifit1, alone or with Ifit2 and Ifit3, was incubated with viral RNA before reverse transcription from a radiolabelled primer specific for the full-length genomic RNA of each virus. cDNA products were resolved by denaturing polyacrylamide gel electrophoresis. Ifit1 was capable of forming a toeprint 6-7 nt downstream of the full-length cDNA product on artificial cap0 RNAs, consistent with binding to the 5’ end (Figure 2C). However, VPg-linked RNAs derived from infected cells were not bound by Ifit1. Addition of Ifit2 and Ifit3 did not affect Ifit1 binding.

### Ifit1 knockout cells have defective innate immune sensing

Ifit1 was previously shown to regulate different stages during innate immune signalling, including signalling downstream of MAVS and TLR3 (27, 29, 30). Therefore, we hypothesised that Ifit1 may mediate its antiviral activity during norovirus infection by promoting the innate immune response to infection. We therefore tested our knockout cell lines for their ability to respond to different stimuli. Wildtype and Ifit1^−/−^ RAW264.7 cells were incubated with LPS or polyI:C, or transfected with polyI:C, to stimulate TLR4, TLR3 or cytoplasmic RNA sensing pathways, respectively. Samples were taken up to 24 hours post infection and RNA or protein was extracted for analysis.

When polyI:C was transfected, to stimulate cytoplasmic RNA sensing, IFNβ expression was strongly upregulated 3 to 9 hours post transfection, decreasing by 12 to 24 hours (Figure 3A). Expression was 4 to 10-fold higher in wild type cells during the peak of expression, compared to Ifit1^−/−^ cells. TNFα was induced to a much lesser extent and expression was comparable between all cell lines (Figure 3B). We observed weak induction of both IFNβ and TNFα when polyI:C was added to the cell culture medium, rather than transfected, and expression levels were comparable between all cell lines tested (Figure 3C-D). This indicates that the differential response in Ifit1^−/−^ cells is specific to cytoplasmic, rather than endosomal, RNA sensing. Cells treated with LPS showed little upregulation of IFNβ mRNA expression when analysed by RT-qPCR (Figure 3E). However, TNFα was strongly upregulated 3 to 6 hours post LPS treatment in all cell lines, returning to near baseline expression by 9 hours post treatment (Figure 3F). At 6 hours, TNFα expression was 2 to 3-fold higher in IFIT1^−/−^ cells compared to wildtype, consistent with a recent report (29).

**Figure 3.**
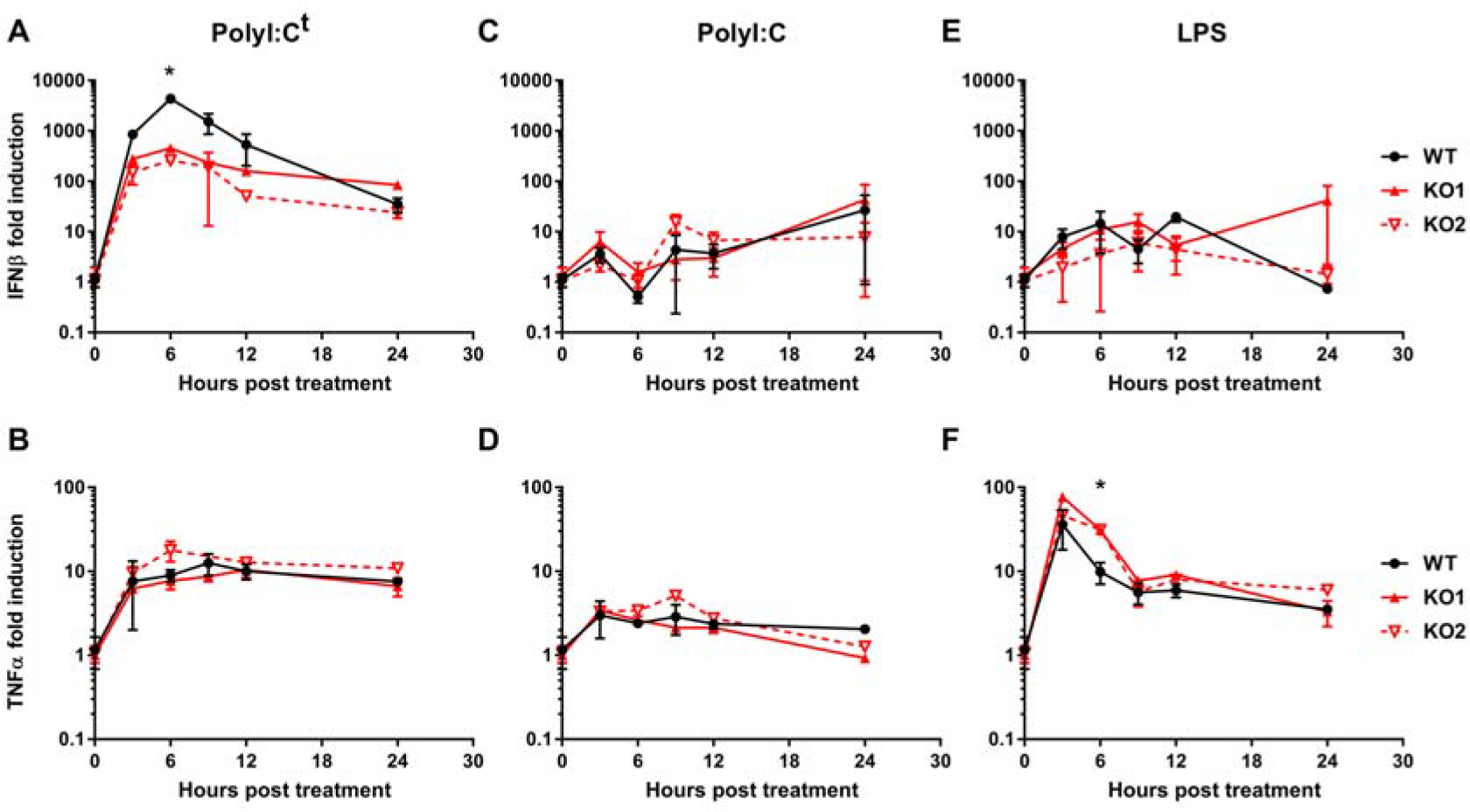
Ifit1 promotes type I IFN expression following cytoplasmic RNA sensing. **A-C.** Wildtype (WT) and Ifit1 knockout (KO) RAW264.7 cells were stimulated with **(A)** 2 μg transfected polyI:C (polyI:C^t^), or **(B)** 1 μg/mL polyI:C or **(C)** 10 ng/mL lipopolysaccharide (LPS) in the cell culture medium. RNA was extracted and analysed by RT-qPCR for IFNβ and TNFα mRNA, expressed as fold induction over untreated cells, normalised against GAPDH (2^−ddCT^). Graphs show the mean and the standard error of three biological replicates. Fold induction was compared between WT and KO cells for each time point by two-tailed student’s t-test. Asterisks indicate that a statistically significant difference (p < 0.05) was observed for both KO cell lines.

## Discussion

Noroviruses replicate in the cytoplasm, where they establish membrane-associated replication complexes in which the viral genome is replicated via a dsRNA intermediate. Cytoplasmic double-stranded RNA sensors, RIG-I and MDA5, are principally responsible for detecting replicating calicivirus RNA, activating the type I IFN response (5–7). This rapid and robust antiviral programme is necessary for viral clearance (4). Here, we demonstrated that the antiviral protein Ifit1 promotes type I IFN responses in RAW264.7 cells and as such contributes to the host antiviral response to restrict murine norovirus infection.

We observed that Ifit1^−/−^ RAW264.7 cells were more susceptible to MNV infection. In most cells, Ifit1 is not expressed to detectable levels under basal conditions, but expression is induced within a few hours of IFN treatment or viral infection (8, 46, 47). As such, we noticed a more pronounced difference between wildtype and Ifit1-deficient cells following a low multiplicity infection, since type I IFN from infected cells will induce naïve cells to establish an antiviral state, including the upregulation of Ifit1 expression.

IFIT proteins have been implicated in regulating different stages of the antiviral and inflammatory responses (reviewed in (48)). In humans, IFIT1 was shown to promote type I IFN expression during alphavirus infection (28). Consistently, a recent study in human and murine macrophages has shown that IFIT1 stimulates type I IFN expression, but represses the inflammatory gene programme, in the acute response following a number of different stimuli (29). The authors suggest that a small population of nuclear IFIT1 can modulate the activity of transcription regulatory complex Sin3A-HDAC2, which is responsible for downregulating both type I IFN and inflammatory gene expression.

Another study has suggested that cytoplasmic IFIT1 downregulates IFN expression by disrupting the MAVS-TBK1-STING signalling axis (27). Together, these studies present a model by which a low level of nuclear IFIT1 promotes type I IFN responses by modulating transcriptional activity. Later in infection, when IFIT1 is highly expressed in the cytoplasm, IFIT1 prevents induction of type IFN I by interfering with MAVS signalling. Consistent with this hypothesis, we observed strong IFNβ expression 3 to 9 hours post stimulation in wild type cells, which sharply decreased from 9 to 24 hours. In Ifit1^−/−^ cells, IFNβ expression was induced to a lesser extent, but remained constant up to 24 hours post stimulation, indicating that Ifit1 may be necessary both to switch on and switch off IFN induction at different stages of the immune response.

In caliciviruses, VPg acts as a substitute for the mRNA 5’ cap, by interacting directly with components of the eIF4F complex, to promote ribosome recruitment via eIF3 (32, 39, 43, 44). Additionally, the VPg of MNV and Norwalk virus, the prototypic strain of human norovirus, may also interact with eIF3 to promote efficient translation initiation (49, 50). IFIT proteins have been reported to interact with eIF3 and inhibit translation initiation on certain mRNA transcripts (22–24). Human IFIT1 binds to the e subunit of eIF3 (22) and can inhibit translation from the hepatitis C virus internal ribosome entry site (IRES) (21). However, IFIT1 cannot inhibit translation from the eIF3-dependent encephalomyocarditis virus IRES (22). Murine Ifit1 and Ifit2 have both been shown to bind to different domains of the eIF3c subunit, causing translation inhibition on luciferase reporter mRNA at micromolar concentrations (46). However, while Ifit3 was also reported to bind to eIF3c, it has no impact on translation (9).

We have demonstrated that 5’ VPg renders calicivirus genomic RNA resistant to Ifit1-mediated translation inhibition. We saw no effect on translation of either capped or VPg-linked RNA when Ifit2 and Ifit3 were added to *in vitro* translation lysates, indicating that neither of these proteins can inhibit translation, despite their potential to interact with eIF3. Therefore, it remains to be determined how IFIT-eIF3 interactions can inhibit translation initiation on some transcripts but not on others.

In summary, despite calicivirus RNA being refractory to translation inhibition by Ifit proteins, we have shown that Ifit1 knockout cells support a higher degree of MNV infection compared to wildtype cells. We observed that Ifit1 promoted type I IFN expression downstream of cytoplasmic dsRNA sensing, suggesting it may play a role in potentiating the host antiviral state. This work contributes to a growing body of evidence that IFIT proteins can modulate innate immune signalling, complementing their role in translation inhibition.

## Data availability

All data underlying the results are available as part of the article and no additional source data are required.

## Competing interests

The authors declare no competing interests.

## Grant Information

This research was funded by a Wellcome Trust/Royal Society Sir Henry Dale Fellowship to TS (202471/Z/16/Z) and a Wellcome Trust Senior Fellowship to IG (207498/Z/17/Z). HM is supported by a Department of Pathology PhD studentship.

